# Multivariate Associations Between Neurophysiological Activity and Sensory Motor Cortical Morphometry in Parkinson’s Disease

**DOI:** 10.1101/2025.07.28.667342

**Authors:** Koorosh Mirpour, Amirreza Alijanpourotaghsara, Nader Pouratian

**Affiliations:** Neurological Surgery Department, UT Southwestern Medical Center, 5303 Harry Hines Blvd, Dallas, 75390, TX, US

**Keywords:** Parkinson’s disease, Deep Brain Stimulation, DBS, Beta Oscillation, Beta Burst, GPi, Motor Cortex, Cortical morphometry, cortical structure, electrophysiological biomarkers, neuroimaging, structural MRI, Gray matter thickness, Gray matter volume, Local field potential, LFP, ECoG

## Abstract

Parkinson’s disease (PD) involves progressive neurodegeneration and distinctive structural and functional alterations in cortico-basal ganglia circuits. This study proposes bridging the gap between structural and functional biomarkers to uncover fundamental mechanisms underlying PD pathophysiology and support more comprehensive diagnostic and therapeutic approaches. We examined 50 PD patients using high-resolution MRI to quantify cortical thickness, surface area, and volume in sensorimotor regions, alongside intraoperative neurophysiological recordings of spectral power, burst parameters, and coherence. Pairwise correlation and Sparse Partial Least Squares analyses revealed significant low-dimensional latent relationships, particularly linking cortical atrophy in Brodmann areas BA1–BA6 with alpha and low-beta burst abnormalities. These associations remained robust after controlling for age, disease duration, and symptom severity, and were absent in a control cohort with essential tremor. These findings highlight the value of multimodal approaches for uncovering structure-function interactions in PD and highlight the potential of integrated biomarkers for improving diagnosis and treatment strategies.

## 1 Introduction

The pathophysiology of Parkinson’s disease (PD) is complex, involving multifaceted changes in brain structure and function. These two usually independent research avenues have significantly advanced our knowledge of PD: electrophysiological studies focusing on neural oscillations and neuroimaging studies investigating brain morphometry.

Electrophysiological investigations have underscored the pivotal role of abnormal neural oscillations within the basal ganglia circuitry of people with PD (PwPD), most notably exaggerated beta-frequency oscillatory activity (13–35 Hz) in the subthalamic nucleus (STN) and globus pallidus internus (GPi) [1, 2]. This heightened beta power is associated with motor symptoms such as rigidity and bradykinesia and decreases during movement preparation and execution [54, 55]. Beta activity is modulated by pharmacological treatments and deep brain stimulation (DBS), with suppression of beta power correlating with improvements in motor function [3, 4]. These findings suggest a direct link between aberrant electrophysiological patterns and the hypodopaminergic state underlying PD symptoms.

The beta band is increasingly subdivided into low beta (13–20 Hz) and high beta (21–35 Hz) frequencies, each likely with unique functional roles [5, 6]. The low beta band, characterized by heightened power within the STN, is closely linked to pathological oscillations and demonstrates greater sensitivity to medication, in contrast to the high beta band, which is less pronounced and less responsive to pharmacological treatments. [3]. Other studies highlight a clear distinction between the local significance of low beta oscillations within the basal ganglia and the broader role of high beta oscillations across the BGTC motor circuit[7]. The same study along with others show that spectral coherence within the cortico-basal ganglia network has emerged as a crucial biomarker for motor control [4, 7, 8]. Abnormal beta bursts—transient increases in beta power—have also been significantly correlated with the severity of motor impairment, offering a more precise explanation of motor deficits than average beta power measurements [9–11].

In parallel, neuroimaging studies have documented progressive structural changes in the brains of PwPD. Cortical thinning has been observed across temporal, parietal, frontal, and occipital regions, particularly in patients with mild cognitive impairment [12, 13]. Reductions in cortical thickness correlate with disease severity, as indicated by Hoehn and Yahr stages and Unified Parkinson’s Disease Rating Scale (UPDRS-III) scores, especially in regions such as the bilateral fusiform gyrus and left temporal pole [14, 15]. Other structural alterations include changes in gray matter (GM) volume [16], cortical microstructure [17], and increased rates of cortical and subcortical atrophy [18]. However, inconsistencies in the literature complicate the interpretation of these findings. While some studies report no significant cortical structural changes in PD [19, 20], others highlight reductions in volume across sensory-motor regions such as the frontal [21, 22] and temporal lobes [23, 24], intraparietal sulcus [25]. These discrepancies may stem from subject heterogeneity, as many studies fail to account for factors like disease duration or clinical subtypes. Moreover, the diverse analytical techniques employed—ranging from region-of-interest approaches to voxel-based morphometry (VBM) and cortical thickness analysis—often yield inconsistent results, limiting their diagnostic sensitivity [26].Despite these advances, the relationship between neurophysiology and structural brain changes in PwPD remains inadequately explored. Sanmartino et al. (2022) [27] proposed that STN LFP beta power might be linked to parallel neurodegenerative processes: one involving GM volume reduction in the dorsal striatum and another concerning cortical thickness reduction in prefrontal “associative” regions. Another recent study by our group examining the relationship between cortical thickness and beta oscillations in PwPD found a significant negative correlation between high-beta (20-35 Hz) power and cortical thickness, suggesting that cortical thinning may contribute to elevated high-beta activity characteristic of PD pathophysiology [28].

While a few studies have documented cortical thinning and increased atrophy in both cortical and subcortical networks [29, 30], the presence of disease-specific regional cortical atrophy as a definitive biomarker for PD remains controversial.

Understanding the interplay between neurophysiology and cortical morphometrics is crucial for gaining insight into the pathophysiology of the cortico-basal ganglia circuitry in PD, especially as oscillations are an emergent property of underlying neural activity which is presumably affected by changes in morphometry. One major obstacle is the multifactorial nature of the disease, involving multiple electrophysiological and structural markers that contribute to a complex network of interactions. Traditional pairwise analyses may be insufficient to capture these comprehensive associations, making it challenging to interpret the underlying pathophysiological mechanisms.

We hypothesize that a general link exists between electrophysiology and cortical morphometrics that extends beyond isolated pairwise correlations between individual biomarkers. Specifically, we propose that dynamic and complex electrophysiological processes—such as bursting activity and coherence—are correlated collectively with structural alterations in the brain. All distributed biomarkers that are extracted from structural alterations are potentially linked to fundamental core pathophysiological mechanisms that are equally associated with the distributed neurophysiology of the disease. While sparse cross-correlation studies may identify incidental relationships between pairs of structural and neurophysiological biomarkers at the face value, they fail to capture the interconnected and holistic nature of these core mechanisms.

## 2 Results

Building on this overarching hypothesis, our aim is to explore the intricate relationship between cerebral morphology and neurophysiology in PD. Specifically, we seek to examine the associations between local cortical thickness, surface area, and volume, and their cross-correlations with neurophysiological biomarkers. To achieve this, we aim to reduce the high-dimensional biomarkers of each modality into lowdimensional latent variables that encapsulate the holistic relationships underlying structure-neurophysiology interactions in PD. Additionally, we investigate a broad spectrum of electrophysiological biomarkers implicated in PD pathophysiology, including beta power subcomponents, coherence, and beta bursts, to determine their collective associations with structural changes in selected sensorimotor cortical regions. To comprehensively uncover these associations, we employ advanced multivariate analytical methods capable of capturing the latent correlations between structural and neurophysiological variables, providing a unified and integrative perspective on the neural correlates of structural alterations in PD.To address these objectives, we employ both traditional pairwise analyses and a more comprehensive examination of associative effects between structural and neurophysiological variables. By utilizing canonical correlation analysis, we aim to maximize the correlation and covariation between latent variables underlying structure and neurophysiology, allowing us to capture the neural correlates of structural alterations in PD beyond single sparse correlations and providing insights into the fundamental relationship between brain structure and function.Our approach recognizes that the variables contributing to the structure-neurophysiology link are not necessarily the same as those correlating with other disease features, such as age, disease duration and behavioral severity scores. By focusing on the associative effects of multiple factors, we strive to unravel the complex interplay between electrophysiological activity and cortical morphometrics in PD. By elucidating the complex interactions between structure and neurophysiology, we hope to lay the ground work to identify novel biomarkers of disease progression and enhance our understanding of the underlying pathophysiological mechanisms in PD.

The study included 50 PwPD, including 13 women with an average age of 64.36 years. Average Unified Parkinson’s Disease Rating Scale (UPDRS) III score was 37.73. Male participants had a mean age of 63.14 years and an average UPDRS score of 39.12, while female participants had a mean age of 67.85 years and an average UPDRS score of 3639. Statistical analyses revealed no significant differences between male and female participants in terms of age (t-test; p=0.062) or UPDRS III scores (t-test; p=0.721) (see Table 1).

**Table 1:**
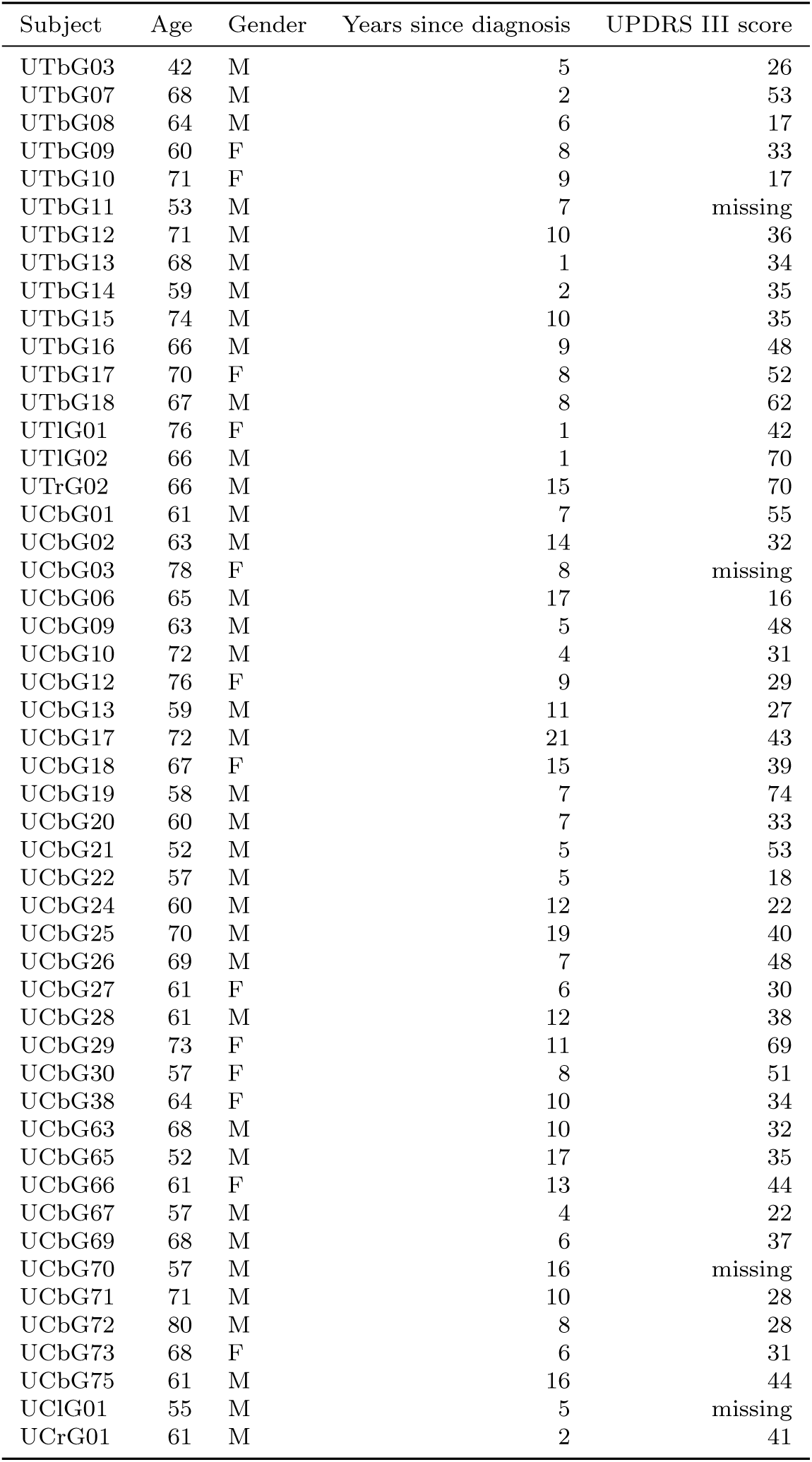
Subject data including Age, Gender, Years since diagnosis, and UPDRS III score.

### 2.1 Pairwise Cross-Correlation Analysis

To explore the relationships between structural and neurophysiological biomarkers, we first calculated Pearson correlation coefficients for all possible pairs of these measures. Figure 1A illustrates these pairwise correlations across the two modalities. To assess the statistical significance of the observed patterns, we performed permutation tests by randomly shuffling the data for each subject and repeating the correlation analysis 1,000 times. Correlations that fell within the extreme 5% of the permutation distribution were considered significant and are plotted in Figure 1B.

**Fig. 1:**
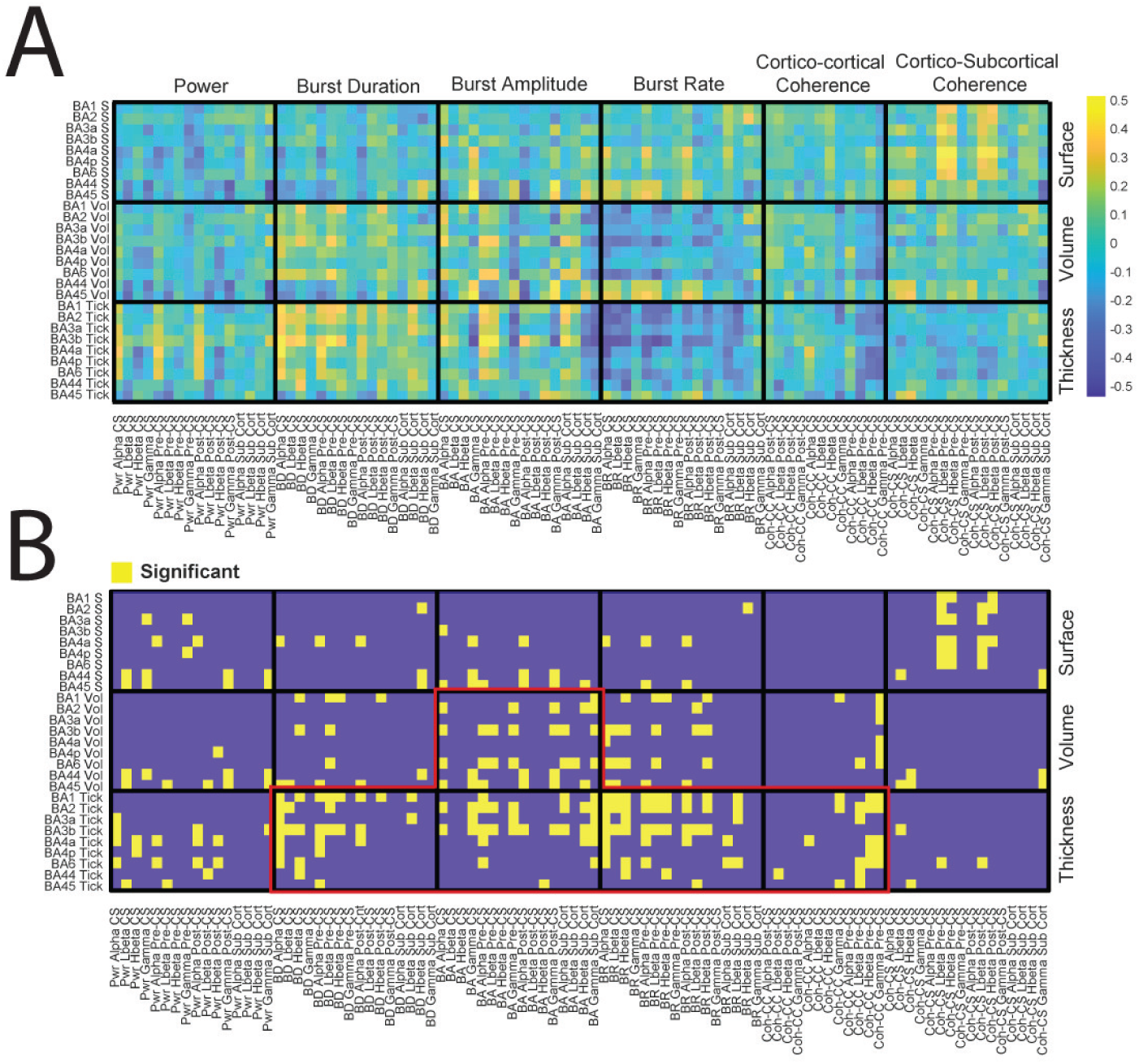
Cross-Correlation Analysis Between Structural and Electrophysiological Measures in PD Patients. (A) Heatmap showing the pairwise Pearson correlation coefficients between structural and electrophysiological measures across 50 PD patients. The structural measures include cortical thickness, volume, and surface area for Brodmann areas (BA1, BA2, BA3a, BA3b, BA4a, BA4p, BA6, BA44, BA45), while the electrophysiological measures encompass power, burst duration, burst amplitude, burst rate, cortico-cortical coherence, and cortico-subcortical coherence, calculated across various frequency bands. The color scale represents the strength and direction of the correlation coefficients.

Although individual pairwise comparisons appear scattered, grouping the variables into broader categories provides clearer insights. In Figure 1, the axes are organized based on morphometric measures (surface area, volume, and cortical thickness on the Y-axis) and neurophysiological measures (power, burst duration, burst amplitude, burst rate, and coherence on the X-axis). Notable patterns are apparent, such as a positive cluster for burst duration and a negative cluster for burst rate (Figure 1A). Permutation tests identified clusters of significant correlations, with red lines indicating boundaries where clusters exceeded three median absolute deviations from the center (Figure 1B).

Applying the Benjamini-Hochberg procedure to control the false discovery rate (*α*=0.05), we highlighted significant correlations marked with asterisks in Figure 2. For example, Figure 2D shows a significant correlation between power and volume and surface area regardless of band. Another example is Figure 2C, it reveals a strong positive correlation between burst duration and cortical thickness, particularly in regions Brodmann areas (BA) 1 to BA6; however, these did not reach statistical significance. Similarly, burst rate exhibited significant correlations with cortical thickness and volume, especially across regions BA1 to BA3b (Figure 2C). All frequency bands demonstrated significant correlations with average cortical thickness (Figure 2F).

**Fig. 2:**
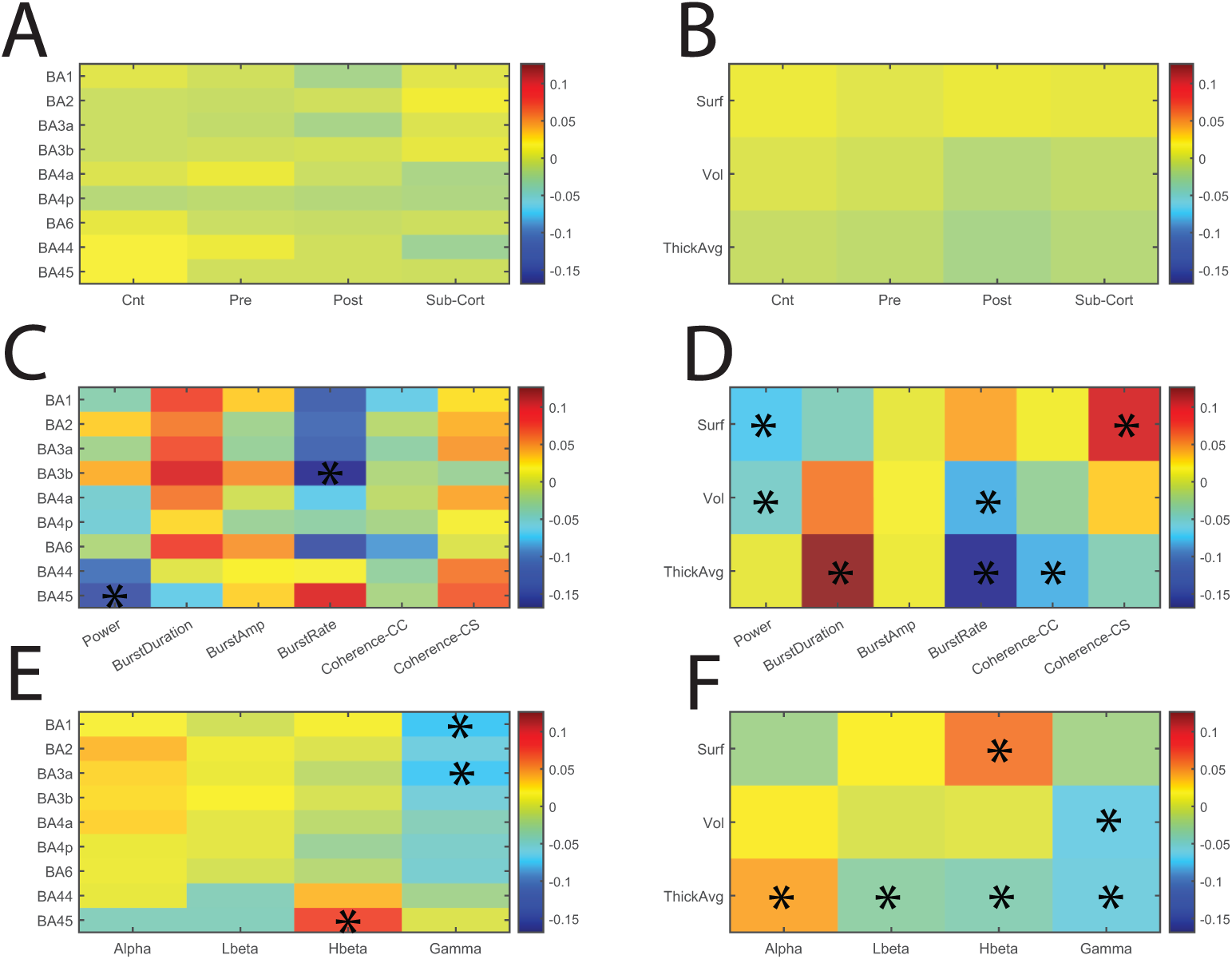
Detailed Analysis of Cross-Correlation Between Structural and Electrophysiological Measures. (A-F) Heatmaps illustrate the pairwise Pearson correlation coefficients between structural and electrophysiological measures, grouped by different subcategories across 50 PD patients. The color scale represents the strength and direction of correlation coefficients. Asterisks (*) indicate significant correlations after permutation testing and False Discovery Rate (FDR) control. (A) Correlation of structural measures (BA1 to BA6) with electrophysiological metrics categorized by electrode location: central (CS), precentral (Pre), postcentral (Post), and subcortical (Sub-Cort). (B) Correlation of surface area (Surf), volume (Vol), and average thickness (Thick Avg) with electrophysiological metrics categorized by electrode location. (C) Correlation of structural measures (BA1 to BA6) with detailed electrophysiological metrics: power, burst duration, burst amplitude, burst rate, cortico-cortical coherence (Coherence-CC), and cortico-subcortical coherence (Coherence-CS). (D) Correlation of surface area, volume, and average thickness with detailed electrophysiological metrics. (E) Correlation of structural measures (BA1 through BA6) with electrophysiological metrics across specific frequency bands: alpha, low beta (Lbeta), high beta (Hbeta), and gamma. (F) Correlation of surface area, volume, and average thickness with electrophysiological metrics across specific frequency bands. BA = Broadman area, Surf = surface area, Vol = volume, Tick Avg= average thickness, LBeta = low beta band, Hbeta = high beta band, CS = central sulcus, Pre-CS = precentral sulcus, Post-CS = postcentral sulcus, Sub-cort = subcortical region (Pallidum), Coherence-CC = cortico-cortical coherence, Coherence-CS = cortico-subcortical coherence. Significant correlations are marked with asterisks in all panels.

Despite revealing significant relationships between certain structural and neurophysiological measures, this pairwise analysis primarily provides a fragmented view of the associations. While clustering schemes were applied to identify patterns, they often showed weak convergence and failed to decisively uncover a fundamental relationship between the two modalities. This limitation underscores the need for more comprehensive methods to capture the interrelated nature of morphometric and neurophysiological variables.

### 2.2 Associative Analysis Using Sparse Partial Least Squares

While pairwise analyses are effective for examining specific relationships between individual biomarkers and morphometric properties, they do not account for the overall associative effects of one modality on the other. To address this limitation, we employed Sparse Partial Least Squares (SPLS) regression to investigate the multivariate associations between structural and neurophysiological variables.

Sparse Partial Least Squares (SPLS) is a multivariate statistical technique designed to identify and quantify relationships between two high-dimensional datasets. By maximizing the covariance between linear combinations of variables from each dataset, SPLS captures the underlying associations while imposing sparsity constraints. These constraints ensure that only the most relevant features contribute to the model, enhancing both interpretability and generalizability. In the context of this study, SPLS enables the discovery of latent dimensions that represent the interconnected dynamics between morphometric and neurophysiological variables, providing a holistic view of their associations. We analyzed the entire dataset using SPLS with a 20% holdout approach across 10 separate data splits. This analysis identified two significant latent dimensions representing associations between structure and neurophysiology. However, only the first latent dimension showed significant correlations between predicted and observed scores in the test sets (training set: *ρ*=0.688, p¡0.001; test set: *ρ*=0.818, p=0.001; Figure 3A). Therefore, our focus is on the first latent dimension.

**Fig. 3:**
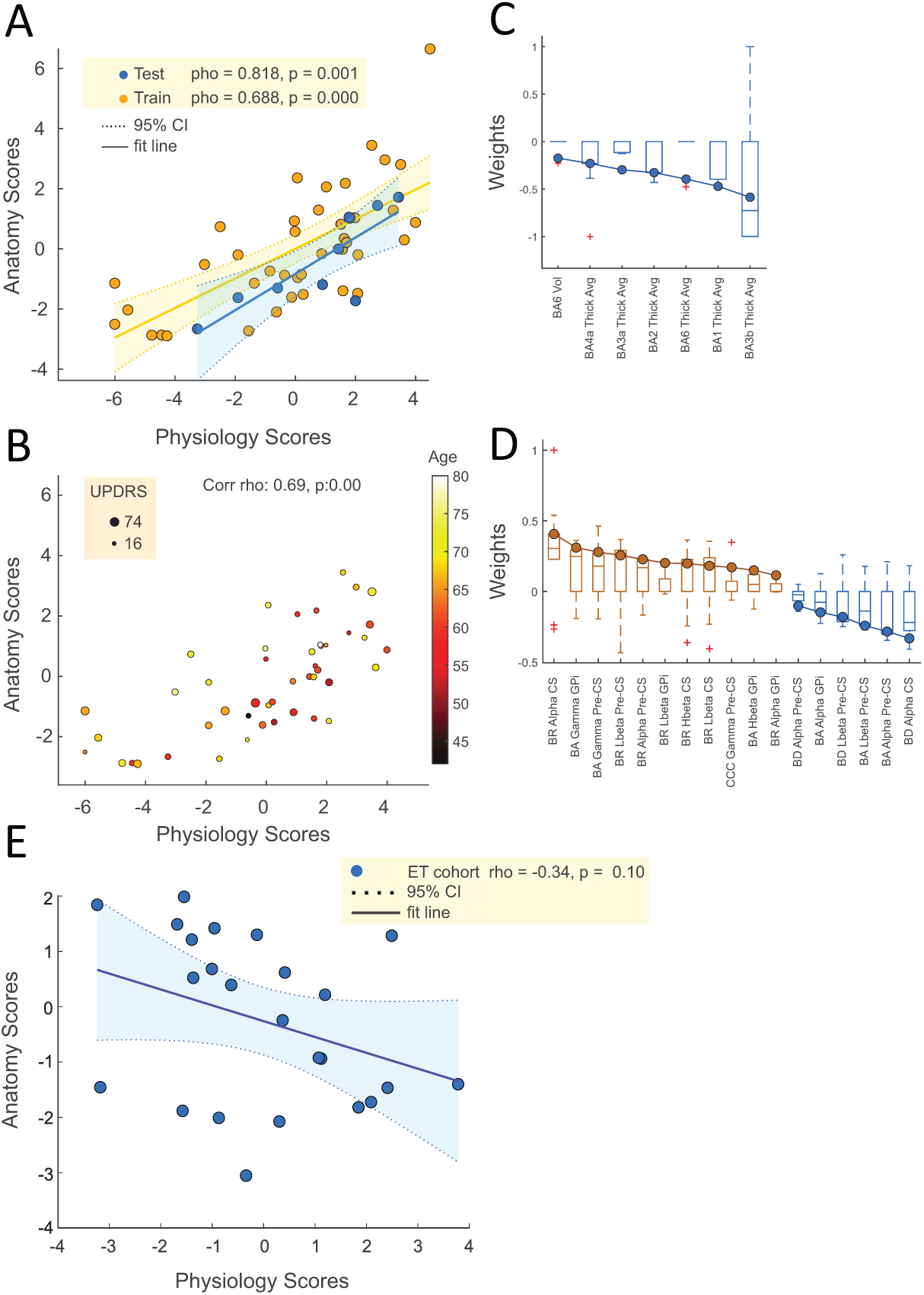
Canonical Correlation Between Structural and Neurophysiological Latent Scores and related weights. (A) Scatterplot illustrating the relationship between structural and neurophysiological latent scores in both the training (orange) and test (blue) sets. The fit line represents the linear regression, and the shaded area indicates the 95% confidence interval (CI). The strong correlation between these scores in both sets (train: *ρ* = 0.688, p ¡ 0.001; test: *ρ* = 0.818, p = 0.001) highlights the robustness of the identified structure-neurophysiology association. (B) Scatterplot showing the relationship between structural and neurophysiological latent scores across all subjects, with color indicating patient age and size representing the Unified Parkinson’s Disease Rating Scale (UPDRS) scores. The correlation coefficient (*ρ* = 0.69, p ¡ 0.001) demonstrates a significant association between the latent scores. (C) Boxplot of the weights for the structural variables contributing to the first latent dimension identified by SPLS. The structural variables include volume (Vol) and average thickness (Thick Avg) for Brodmann areas (BA) 6, 4a, 3a, 3b, 2, and 1. The negative weights indicate a strong negative association between cortical thickness and the neurophysiology latent dimension, particularly in the sensorimotor regions BA3b, BA1, BA6, BA2, and BA4a, suggesting that thinning in these regions is strongly linked to electrophysiological alterations in PD. (D) Boxplot of the weights for the neurophysiological variables contributing to the first latent dimension. The neurophysiological variables include burst rate (BR), burst duration (BD), and cortico-cortical coherence (CCC) across different frequency bands (alpha, low beta [Lbeta], high beta [Hbeta], gamma) and locations (central [CS], precentral [Pre-CS], GPi). Positive weights indicate a positive association with the structure’s latent dimension, with central alpha burst rate and gamma burst amplitude showing the strongest contributions. The negative weights, particularly in low beta burst duration and alpha burst duration, highlight their negative association with cortical thickness. (E) Scatterplot illustrating the relationship between structural and neurophysiological latent scores in ET control cohort. The fit line represents the linear regression, and the shaded area indicates the 95% confidence interval (CI). No significant correlation was found between structural and neurophysiological latent scores (*ρ* = -0.342, p = 0.102). BA = Broadman area, Vol = volume, Tick Avg = average thickness, LBeta = low beta band, Hbeta = high beta band, CS = central sulcus, Pre-CS = precentral sulcus, Post-CS = postcentral sulcus.

Table 2 summarizes the outcomes for the first latent dimension across all 10 splits. Five splits (bolded in Table 2) yielded significant results in the test samples that met the criteria for passing the omnibus hypothesis (Nichols and Holmes, 2001). To ensure robustness, we present results from the split that demonstrated the optimal balance of generalizability (assessed by out-of-sample correlation on the holdout set) and stability (assessed by the consistency of weights across optimization sets).

**Table 2:**
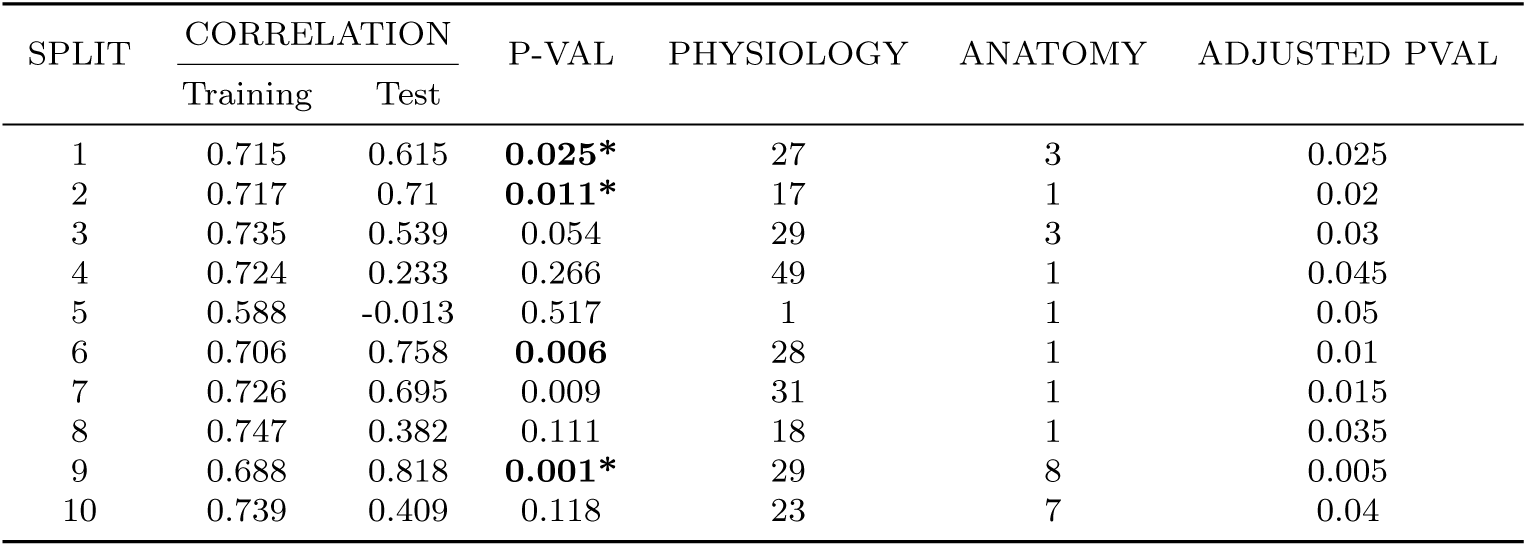
Summary of Sparse Partial Least Squares (SPLS) Model Performance Across Different Data Splits. The table presents the results of the SPLS analysis across 10 different data splits. For each split, the training and test set correlations are shown, alongside the corresponding test p-values, the number of selected physiology features, and the number of selected anatomy features. The final column lists the Benjamini-Hochberg (BH) adjusted p-values for significance. Significant correlations between anatomy and physiology latent dimensions are highlighted in bold. The results indicate the robustness and generalizability of the model, with certain splits (e.g., 2, 6, 7, and 9) demonstrating particularly strong and significant associations. * Significant with BH adjusted p-values.

| SPLIT | CORRELATION |  | P-VAL | PHYSIOLOGY | ANATOMY | ADJUSTED PVAL |
| --- | --- | --- | --- | --- | --- | --- |
|  | Training | Test |  |  |  |  |
| 1 | 0.715 | 0.615 | **0.025\*** | 27 | 3 | 0.025 |
| 2 | 0.717 | 0.71 | **0.011\*** | 17 | 1 | 0.02 |
| 3 | 0.735 | 0.539 | 0.054 | 29 | 3 | 0.03 |
| 4 | 0.724 | 0.233 | 0.266 | 49 | 1 | 0.045 |
| 5 | 0.588 | -0.013 | 0.517 | 1 | 1 | 0.05 |
| 6 | 0.706 | 0.758 | **0.006** | 28 | 1 | 0.01 |
| 7 | 0.726 | 0.695 | 0.009 | 31 | 1 | 0.015 |
| 8 | 0.747 | 0.382 | 0.111 | 18 | 1 | 0.035 |
| 9 | 0.688 | 0.818 | **0.001\*** | 29 | 8 | 0.005 |
| 10 | 0.739 | 0.409 | 0.118 | 23 | 7 | 0.04 |

In this optimal split (test set: *ρ*=0.688, p=0.001; Table 2, row 9), the SPLS model revealed a strong negative correlation with average cortical thickness in regions BA3b, BA1, BA6, BA2, BA3a, and BA4a (Figure 3C). The most influential positive neurophysiological contributors were central alpha burst rate, globus pallidus internus (GPi) gamma burst amplitude, and precentral gamma burst amplitude. Conversely, the strongest negative associations were observed with central alpha burst duration, precentral alpha burst amplitude, and precentral low beta burst amplitude (Figure 3D).

Figure 3A illustrates the correlation between the morphometric and neurophysiology latent variables in both the training and test sets for this optimal split. The high degree of alignment demonstrates the effectiveness of our cross-validation and regularization strategies.Consistent with SPLS methodology, the model produced sparse weight vectors, selecting between 3 and 51 features out of 92 possible neurophysiological variables (mean selection rate: 27.4%±1.4% SEM) and between 1 and 4 out of 36 possible morphometric variables (mean selection rate: 10.0%±1.9% SEM) across different splits. Notably, the best-performing models utilized a smaller number of features, underscoring the importance of feature selection for enhancing interpretability and generalizability.

The weight vectors for the significant neurophysiological measures (absolute weights *>* 0.1) are displayed in Figure 3D. These measures were the most influential contributors to constructing the latent variables, ultimately driving the observed correlations between the two modalities. For structural measures, the average cortical thickness in regions BA3b, BA1, BA6, BA2, BA3a, BA4a, and the volume of BA6 showed significant negative associations (Figure 3C). The optimal model included 29 non-zero weights out of 92 neurophysiological variables, primarily related to burst duration and amplitude within the alpha, gamma, and low beta bands. Central alpha burst duration and central alpha burst rate were the most influential negative and positive weights, respectively (Figure 3D).

Although the second structure-neurophysiology latent space model reached statistical significance (best p=0.036, *ρ*=0.609), it did not reveal consistent sets of weights across the 10 splits and was not significant after Benjamini-Hochberg adjustment. Therefore, we do not present detailed findings for this model.

Scatterplots of the structure and neurophysiology latent scores, derived from applying the weights of the optimal model to all data points, visualize how the structure-neurophysiology relationship manifests across the latent space (Figure 3). The first latent variable showed a Pearson correlation coefficient of 0.649 (p¡0.001). In Figure 3B, the color and size of the data points represent the patients’ age and UPDRS scores, respectively. Subtle patterns suggest potential correlations between disease severity, age, and the latent space correlation effect.

### 2.3 Assessment of Age and Disease Severity and Duration Effects

To investigate the potential influence of age and disease severity on the observed structure-neurophysiology relationship, we conducted partial correlation analyses. The results of partial correlation analysis between structure latent scores, neurophysiology latent scores, UPDRS scores, years since diagnosis and age are presented in Table 3. The correlation between neurophysiological and structural scores is strong and significant (*ρ*= 0.701, p ¡ 0.001) while controlling for age, years since diagnosis (as a measure for disease duration), and UPDRS scores. This indicates that the observed canonical relationship between latent scores cannot be solely attributed to factors such as age or the chronic effects of disease progression.

**Table 3:** Summary of Sparse Partial Least Squares (SPLS) Model Performance Across Different Data Splits. The table presents the results of the SPLS analysis across ten different data splits. For each split, the training and test set correlations are shown alongside the corresponding test p-values, the number of selected physiology features, and the number of selected anatomy features. The final column lists the Benjamini-Hochberg (BH) adjusted p-values for significance. Significant correlations between anatomy and physiology latent dimensions are highlighted in bold. The results indicate the robustness and generalizability of the model, with certain splits (e.g., 2, 6, 7, and 9) demonstrating particularly strong and significant associations. * Significant with BH adjusted p-values.

|  | Physiology | Anatomy | UPDRS score | Age |
| --- | --- | --- | --- | --- |
| Physiology | - | - | - | - |
| Anatomy | **0.701 (p << 0.001)\*** | - | - | - |
| UPDRS score | -0.182 (p = 0.244) | -0.003 (p = 0.984) | - | - |
| Age | -0.292 (p = 0.057) | **0.327 (p = 0.032)\*** | 0.022 (p = 0.887) | - |
| Years since diagnosis | **0.382 (p = 0.012)\*** | -0.093 (p = 0.554) | 0.216 (p = 0.165) | 0.092 (p = 0.557) |

It is noteworthy that we did not initially predefine PD-specific morphometric or electrophysiological biomarkers to include in the SPLS models. Therefore, while the strong correlation between latent scores is not entirely driven by chronic disease changes or age according to partial correlation analysis, these scores likely reflect PD-related changes as well as structure-neurophysiology correlations influenced by other factors. This hypothesis is supported by the lack of significant correlations between UPDRS scores and either structure or neurophysiology latent scores (*ρ*= -0.188, - 0.139; p = 0.212, 0.356, respectively). We propose that the latent scores are influenced by a combination of variables, some directly related to disease severity and others independent of it.

To further disentangle these two sets of variables, we performed stepwise multiple regression analyses to identify factors linked to UPDRS scores. Using structure and neurophysiology biomarkers weighted by split weights, we applied a stepwise approach optimized with the Akaike Information Criterion (AIC) to select the best predictors. Both models converged on a small subset of variables, revealing significant correlations between UPDRS scores and specific biomarkers (r² = 0.227, p = 0.004 for neurophysiology; r² = 0.142, p = 0.037 for structure). The results of the stepwise regression analyses are highlighted in Tables 4 and 5, where variables identified as PD-related biomarkers are indicated.

**Table 4:**
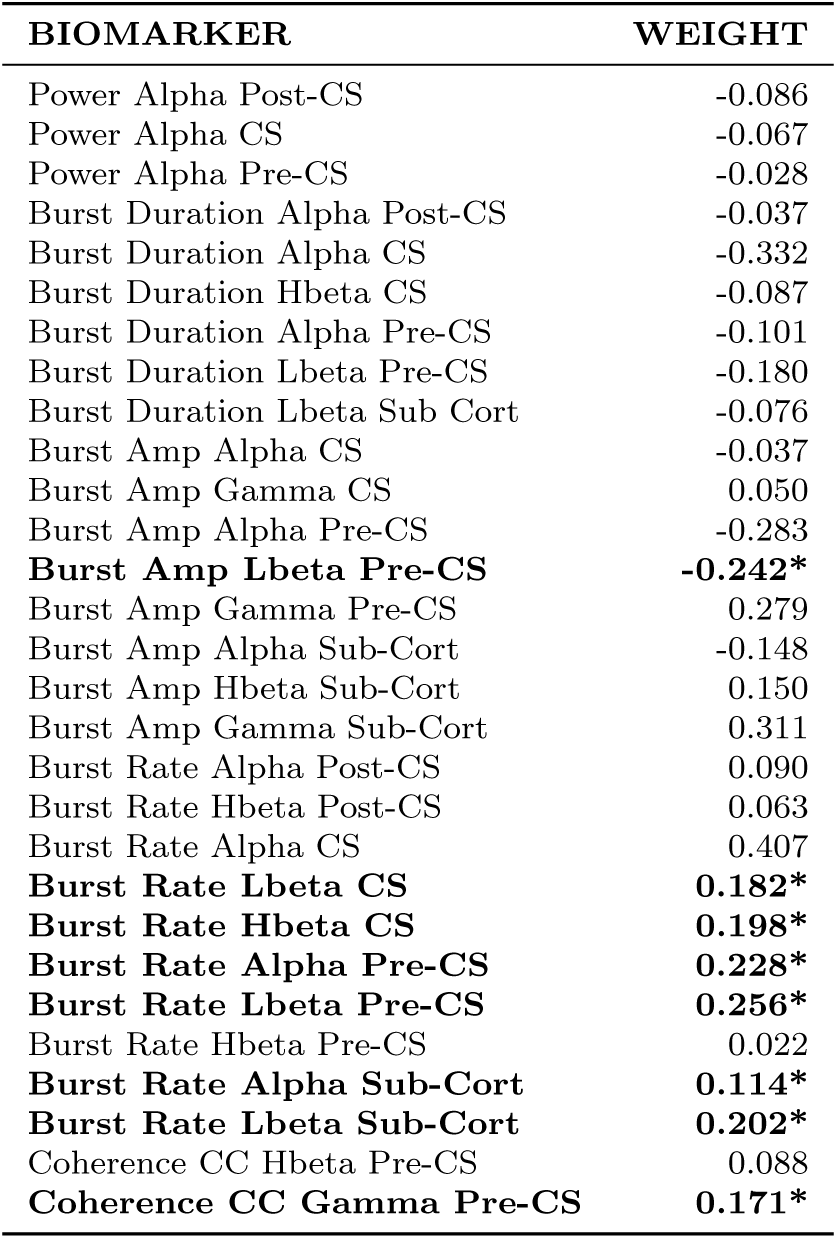
SPLS electrophysiology weights. Nonzero Weights of SPLS electrophysiology. The bold rows are factors that have shown a significant correlation with UPDRS III scores using stepwise linear regression. Abbreviations: Pre-CS precentral sulcus, Post-CS is postcentral sulcus, Sub-Cort is subcortical, Lbeta is low beta, Lbeta is high beta, CC is cortico-cortical. * Significant with BH adjusted p-values.

| BIOMARKER | WEIGHT |
| --- | --- |
| Power Alpha Post-CS | -0.086 |
| Power Alpha CS | -0.067 |
| Power Alpha Pre-CS | -0.028 |
| Burst Duration Alpha Post-CS | -0.037 |
| Burst Duration Alpha CS | -0.332 |
| Burst Duration Hbeta CS | -0.087 |
| Burst Duration Alpha Pre-CS | -0.101 |
| Burst Duration Lbeta Pre-CS | -0.180 |
| Burst Duration Lbeta Sub Cort | -0.076 |
| Burst Amp Alpha CS | -0.037 |
| Burst Amp Gamma CS | 0.050 |
| Burst Amp Alpha Pre-CS | -0.283 |
| **Burst Amp Lbeta Pre-CS** | **-0.242\*** |
| Burst Amp Gamma Pre-CS | 0.279 |
| Burst Amp Alpha Sub-Cort | -0.148 |
| Burst Amp Hbeta Sub-Cort | 0.150 |
| Burst Amp Gamma Sub-Cort | 0.311 |
| Burst Rate Alpha Post-CS | 0.090 |
| Burst Rate Hbeta Post-CS | 0.063 |
| Burst Rate Alpha CS | 0.407 |
| **Burst Rate Lbeta CS** | **0.182\*** |
| **Burst Rate Hbeta CS** | **0.198\*** |
| **Burst Rate Alpha Pre-CS** | **0.228\*** |
| **Burst Rate Lbeta Pre-CS** | **0.256\*** |
| Burst Rate Hbeta Pre-CS | 0.022 |
| **Burst Rate Alpha Sub-Cort** | **0.114\*** |
| **Burst Rate Lbeta Sub-Cort** | **0.202\*** |
| Coherence CC Hbeta Pre-CS | 0.088 |
| **Coherence CC Gamma Pre-CS** | **0.171\*** |

**Table 5:** SPLS anatomy weights. All nonzero Weights of SPLS Anatomy. The bold rows are factors that have shown a significant correlation with UPDRS III scores using stepwise linear regression. Abbreviations: Thick is thickness, Avg is average.

| BIOMARKER | WEIGHT |
| --- | --- |
| BA3b Gray Vol | -0.017552 |
| BA6 Gray Vol | -0.17451 |
| BA1 Thick Avg | -0.47115 |
| BA2 Thick Avg | -0.32683 |
| BA3a Thick Avg | -0.29708 |
| BA3b Thick Avg | -0.58596 |
| BA4a Thick Avg | -0.2302 |
| BA6 Thick Avg | -0.39477 |

Finally, to evaluate whether the model and latent space were specifically related to PD pathophysiology, we tested the approach on a cohort of 24 essential tremor (ET) patients. Morphometric and electrophysiological variables were calculated for the ET cohort, and the SPLS weights from the best-performing split reported in this study were applied to compute the latent space for these patients. The resulting correlation between structural and neurophysiological latent scores in the ET cohort did not show a significant association (Figure 3E, *ρ* = -0.342, p = 0.102). This control analysis suggests that the model and its identified latent dimensions are specifically tuned to the pathophysiological mechanisms underlying PD, as it failed to demonstrate similar correlations in a non-PD cohort.

## 3 Discussion

### 3.1 Multivariate and Pairwise Analyses Reveal Structural-Functional Associations

Utilizing both traditional pairwise analyses and advanced multivariate methods, we uncovered an underlying relationship that extends beyond isolated pairwise correlations between cortical morphometry and neurophysiological variables within the motor cortico-basal ganglia network that is specific to PwPD.

Pairwise cross-correlation analysis revealed significant relationships between specific electrophysiological measures and structural features. Notably, prolonged burst durations and cortical-subcortical coherence were positively correlated with cortical thickness and surface area, suggesting that differences in cortical morphometry may contribute to changes in oscillatory dynamics. Conversely, increased burst rates and cortico-cortical coherence, specifically between the primary motor and either premotor or primary somatosensory cortex, were negatively correlated with cortical thickness and volume. This negative correlation could reflect two seemingly paradoxical scenarios: increased coherence despite physical shrinkage, or decreased coherence with relative physical expansion. In both cases, a compensatory mechanism becomes plausible. When cortical volume or thickness decreases, the remaining pathways may enhance coherence or signal dynamics to maintain information flow. Conversely, in scenarios with relative increases in cortico-cortical pathways, a reduction in coherence may serve to regulate synchronized activity and prevent excessive oscillation.

### 3.2 Compensatory Mechanisms and Latent Multivariate Associations

While there remains a logical leap in directly attributing this pattern to compensation, it is reasonable to hypothesize that coherence and signal dynamics (represented here by bursting) adjustments may serve as a compensatory mechanism, as these can be more readily modulated by altering the firing rates of neurons. These relationships were particularly evident in sensorimotor regions corresponding to Brodmann areas BA1 to BA6. Across all frequency bands, significant correlations were observed between electrophysiological biomarkers and average cortical thickness (Figure 2D), indicating widespread interactions between electrophysiological and structural variables. The associative analysis using Sparse Partial Least Squares (SPLS) further elucidated these relationships by identifying significant latent dimensions that capture the multivariate associations between electrophysiological and structural variables. The strongest associative effects were associated with low beta, alpha and gamma oscillatory activity in precentral, central, and subcortical regions, the globus pallidus internus (GPi). This supports the hypothesis that abnormal alpha and beta activity reflects underlying structural degeneration. To validate this hypothesis, we applied the model to an essential tremor (ET) cohort, which failed to demonstrate the same level of significant correlations, further reinforcing the specificity of the observed associations in PD.

### 3.3 Regional Contributions and Disease Progression

The analysis also identified BA1 and BA3b thickness as key contributors to the observed associations, suggesting that early changes in these regions may play a role in perpetuating the cycle of electrophysiological abnormalities and motor symptoms in PD. Notably, our partial correlation analysis revealed that disease duration was correlated with latent electrophysiology scores but not with structure scores. While this might seem counterintuitive, it implies that structural changes do not progress gradually throughout all stages of the disease. This observation aligns with previous structural studies showing that PD-related cortical thinning occurs more rapidly during the early stages even before clinical manifestations [31, 32]. These findings suggest that early structural changes in regions like BA1 and BA3b may set the stage for ongoing electrophysiological disruptions, perpetuating the pathophysiological processes underlying PD symptoms [14, 18, 29, 33].

Our findings remained significant after controlling for potential confounding factors such as age, disease duration, and disease severity as measured by UPDRS scores. This indicates that the observed correlations are more likely to be reflective of disease-specific pathophysiological processes rather than being artifacts of generalized aging, the chronic effects of prolonged illness, or the accumulated impact of years of motor impairment. Furthermore, the lack of correlation between the magnitude of the structure-neurophysiology link and variables like age, disease duration, and UPDRS scores suggests that the factors driving this relationship are distinct from conventional biomarkers typically associated with disease severity and progression. This divergence highlights the potential for uncovering novel biomarkers and previously unexplored aspects of PD pathophysiology. However, further fundamental research is required to fully elucidate the mechanistic relationship between these conventional biomarkers and the structure-neurophysiology association we have identified. This could pave the way for a deeper understanding of PD and the development of new diagnostic and therapeutic strategies. The consistency of these findings across multiple data splits and cross-validation procedures further underscores their validity and potential clinical relevance.

### 3.4 Structural-Functional Integration and Biomarker Specificity

Our results strongly support the hypothesis that dynamic, complex electrophysiological factors are closely correlated with structural brain alterations in PD. Specifically, burst duration in motor, premotor, and pallidal regions, as well as burst rates (Table 4), combined with cortical volume and average thickness (Table 5) in the structural modality, indicate a fundamental relationship between brain structure and electrophysiology. This relationship likely reflects core mechanisms underpinning PD pathophysiology. While other studies have captured similar underlying mechanisms using diverse methods and modalities, differences in methodologies and measures make direct, one-to-one comparisons challenging, underscoring the existence of such fundamental principles rather than specific metrics [34, 35].

Our further analysis using stepwise linear regression highlights that only a subset of these biomarkers—namely alpha and beta range burst rates in central, precentral, and pallidal regions, along with cortical thinning in the somatosensory and premotor cortices—were significantly linked to PD severity. This finding suggests that while both disease-related and other biomarkers contribute to the overall latent variables, the identified subset may be particularly relevant for disease progression and symptom manifestation.

### 3.5 Oscillatory Dynamics, Regional Specificity, and Clinical Implications

Importantly, the convergence of structural and functional biomarkers into shared highly correlated latent variables highlights a key observation: stationary structural changes serve as a foundational substrate for the dynamic alterations observed in electrophysiological activity. In the context of PD, atrophy of specific cortical regions may disrupt normal oscillatory patterns, particularly in the alpha and beta bands, leading to the emergence of abnormal and compensatory oscillatory activity. For instance, the negative correlation between burst rates and cortical thickness suggests that as structural integrity declines, remaining pathways may enhance coherence or burst rates to maintain neural communication and motor function. We propose that compensatory mechanisms are particularly likely during the early stages of PD [31, 32, 36], since structural alterations are present but motor signs and symptoms have yet to manifest. This compensatory adjustment highlights the intricate balance between structural degeneration and functional adaptation in PD [35, 37, 38].

The involvement of alpha and low beta oscillations in precentral, central, and subcortical regions further underscores the complexity of electrophysiological changes in PD. Alpha oscillations are crucial for sensorimotor processing and cortical excitability [5, 39],while beta oscillations are strongly associated with motor impairments in PD [2, 7, 8, 40]. Alterations in burst amplitude and duration within these frequency bands not only contribute to the electrophysiological abnormalities observed in PD [10, 11] but also appear to be reflected in cortical morphometric changes.

The identification of BA3b volume and thickness as key contributors highlights the critical role of the primary somatosensory cortex in PD. Changes in this region may disrupt sensory feedback mechanisms that are essential for motor control, thereby perpetuating electrophysiological abnormalities and motor symptoms [41]. Notably, our partial correlation analysis revealed that disease duration was associated with electrophysiological latent scores but not with structural latent scores. Although this may seem counterintuitive, it aligns with evidence suggesting that cortical atrophy occurs more rapidly in the early stages of PD. This pattern suggests that structural changes in regions like BA3b may be early drivers of network-level disruptions, with downstream effects on oscillatory activity and motor symptoms [42].

Moreover, our results align with previous neuroimaging studies reporting cortical thinning in sensorimotor areas of PD patients [13, 43]. The observed associations between cortical thinning and electrophysiological alterations provide a potential explanation for the structural-functional disconnects [35] often observed in PD. Specifically, structural atrophy in regions integral to motor control may impair the transmission of neural signals, leading to altered oscillatory dynamics, changes in coherence patterns, and functional network reorganization. These findings emphasize the importance of considering both structural degeneration and electrophysiological disruptions as interconnected processes in the pathophysiology of PD. Our results are consistent with studies suggesting that combined structural and network-level metrics offer critical insights into PD-related network reorganization [44].

By focusing on cortical thickness and a comprehensive range of electrophysiological biomarkers, our study extends previous findings and highlights the role of sensorimotor regions as key areas of structural-functional integration in PD. Understanding the link between electrophysiological biomarkers and cortical morphometric changes has significant clinical implications. Robust associations between structural and functional biomarkers could serve as a foundation for developing comprehensive diagnostic tools and improving disease staging and progression monitoring. Structural measures that strongly correlate with electrophysiology, such as cortical thinning in specific regions, may also serve as biomarkers for the early detection and assessment of PD severity.

These findings suggest that PD progression is driven not only by dopaminergic deficits but also by concurrent structural and electrophysiological changes [45]. Modulating specific oscillatory activities through pharmacological treatments or noninvasive brain stimulation techniques may influence cortical structural changes and, consequently, disease progression. A multimodal approach that integrates neuroimaging and electrophysiology holds great promise for improving the diagnosis, monitoring, and treatment of PD. Such an approach could facilitate personalized therapeutic strategies aimed at mitigating both structural degeneration and abnormal oscillatory activity, ultimately improving patient outcomes and quality of life.

### 3.6 Limitations and Future Directions

The cross-sectional design of this study precludes causal inferences about the relationship between electrophysiological alterations and morphometric differences. Longitudinal studies are needed to determine whether these markers can predict disease progression and response to treatment, potentially paving the way for more personalized and effective therapeutic strategies in PD management.

The challenges associated with collecting electrophysiological data during deep brain stimulation (DBS) surgery limited the number of cases and regions included in the study, which consequently restricted the range of variables analyzed. We chose to focus on cortical structural measures, particularly within the sensory-motor cortex, as these regions have been relatively less studied in terms of their structural correlations with electrophysiology and were a central focus of our research. In contrast, subcortical structures such as the pallidum and STN require more precise structural methods for segmentation and volumetric analysis to ensure accurate correlation with electrophysiological data. Investigating these subcortical gray nuclei will require a dedicated study utilizing advanced volumetric and shape analysis techniques, including subregional breakdowns, to provide a more detailed and focused understanding of their role in PD pathophysiology. Additionally, the sample size, although adequate for SPLS analysis, limits the generalizability of the findings. Larger, multicenter studies could enhance the robustness of the results and allow for subgroup analyses, such as exploring differences based on disease subtypes, treatment regimens, or genetic factors.

## 4 Conclusion

There are complex interactions between cortical structure and electrophysiological dynamics in PwPD. The significant associations between specific electrophysiological biomarkers and cortical thinning in sensorimotor regions underscore the interconnected nature of structural and functional alterations in PD.

Overall, the integration of structural and functional data presents a promising avenue for advancing our understanding of PD and gaining greater insights into pathoetiologic mechanisms of disease. Future research should focus on longitudinal studies to determine whether these markers can predict disease progression and response to treatment, thereby enhancing our ability to diagnose and monitor PD progression and optimize therapeutic interventions. The potential to develop more personalized and effective therapeutic strategies could significantly impact the management of PD, ultimately improving the quality of life for patients.

## 5 Methods

### 5.1 Participants and Ethical Approval

This study was approved by the Institutional Review Boards of both the University of California, Los Angeles (UCLA), and UT Southwestern Medical Center at Dallas. All participants provided written informed consent in accordance with the Declaration of Helsinki. Deep Brain Stimulation (DBS) implantations and target selection were based solely on clinical grounds and indication, based on recommendations of an interdisciplinary clinical team. Data was derived from 50 patients diagnosed with PD who underwent DBS implantation surgery. Bilateral GPi leads were implanted in all but five patients, who received unilateral implantations (3 left, and 2 right). Detailed patient demographics are provided in Table 1. To adhere to standard clinical protocols, all patients presented for surgery in a practical off-state, with PD medications withheld for at least 12 hours prior to the start of surgery.

### 5.2 Surgical Procedure and Electrophysiological Recordings

Following preoperative clinical stereotactic planning, burr holes were created for DBS lead placement. A subdural electrocorticography (ECoG) strip with eight contacts (4 mm platinum-iridium electrodes spaced 1 cm apart; AdTech Medical) was inserted subdurally in a posterior direction through the burr hole. Once a clinical baseline was established after discontinuation of propofol, DBS leads were implanted using microelectrode recordings, kinesthetic mapping, and macroelectrode stimulation testing. Experimental recordings from the deep brain leads and ECoGs (Figure 4A) were performed simultaneously while patients were awake, at least 30 minutes after cessation of propofol sedation.

**Fig. 4:**
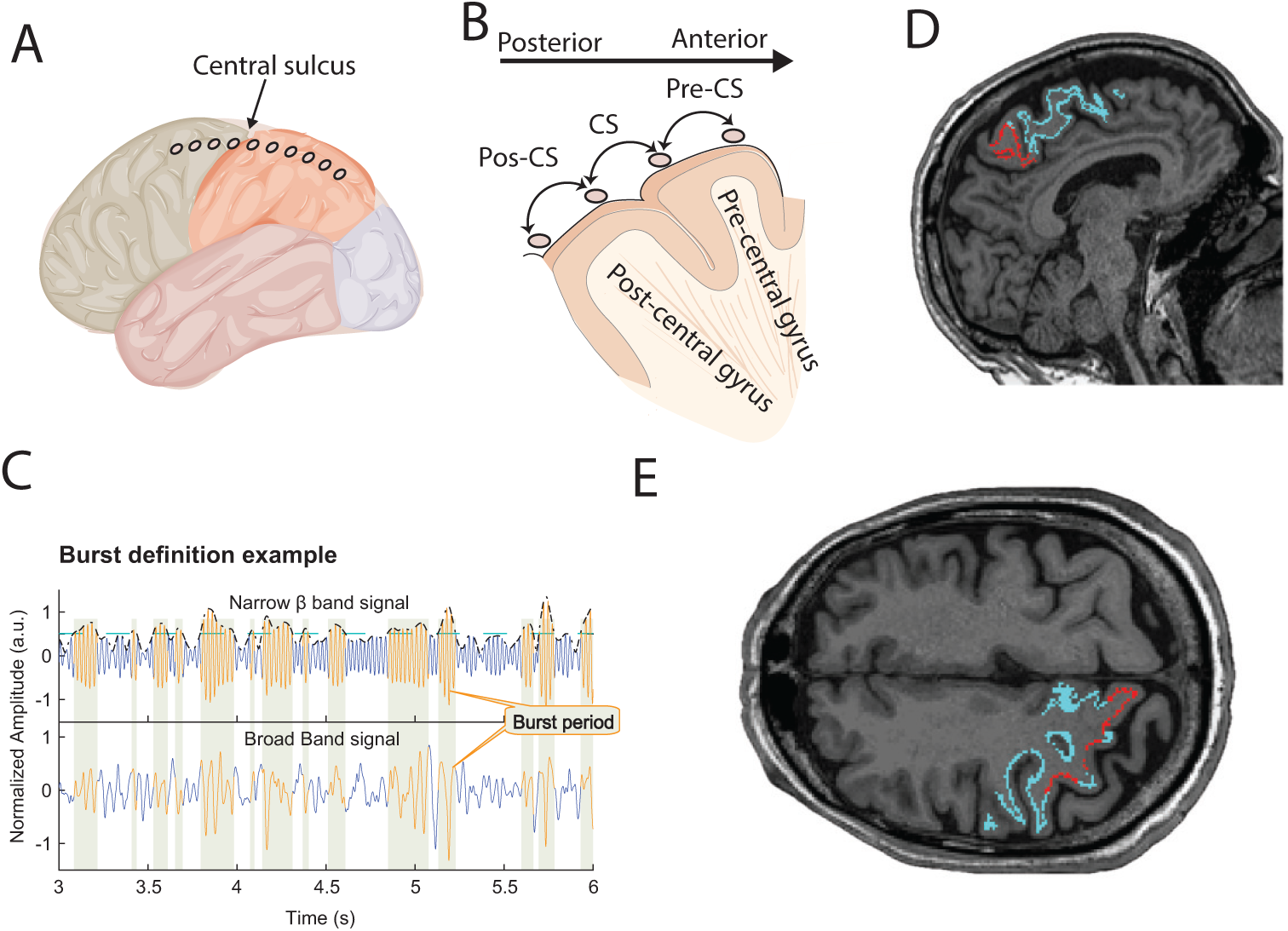
Cross-Correlation Analysis Between Structural and Electrophysiological Measures in PD Patients. (A) Heatmap showing the pairwise Pearson correlation coefficients between structural and electrophysiological measures across 50 PD patients. The structural measures include cortical thickness, volume, and surface area for Brodmann areas (BA1, BA2, BA3a, BA3b, BA4a, BA4p, BA6, BA44, BA45), while the electrophysiological measures encompass power, burst duration, burst amplitude, burst rate, cortico-cortical coherence, and cortico-subcortical coherence, calculated across various frequency bands. The color scale represents the strength and direction of the correlation coefficients.

Neurophysiological recordings were conducted at a sampling rate of 2,400 Hz at UCLA and 1,375 Hz at UT Southwestern (UTSW), with a bandpass filter set between 0.1 and 1,000 Hz. At UCLA, recordings utilized the BCI2000 v3 system connected to amplifiers (g.Tec, g.USBamp 2.0), and at UTSW, the NeuroOmega system was used, with a single scalp reference electrode. Electrode locations were determined postoperatively through a combination of preoperative magnetic resonance imaging (MRI), intraoperative fluoroscopy, and postoperative CT scans, following established protocols [2, 7, 46] . The electrode contact located anterior to the central sulcus was designated as overlying the precentral sulcus (primary motor cortex), while the contact immediately anterior to this was identified as corresponding to the premotor cortex (Figure 1B).

### 5.3 Imaging Acquisition and Processing

High-resolution three-dimensional pre-contrast T1-weighted structural MRI images were acquired for each participant. Imaging sequences included standard T1-weighted (T1W), T2-weighted (T2W), and either focused gradient-echo acquisition for deep brain structures (FGATIR) or subthalamic nucleus (STN)-targeted T2 imaging. The specific sequences and scanner types (Siemens Skyra or Philips Ingenia) varied depending on the center and imaging protocol.

Structural segmentation and processing were performed using FreeSurfer (version 7.3.2; Fischl, 2012), which included skull stripping, normalization, cortical and subcortical segmentation, surface generation, and full cortical reconstruction. We focused on the thickness and volume of sensorimotor areas around the central sulcus, specifically nine Brodmann areas (BAs) segmented by FreeSurfer: somatosensory areas (BA1, BA2, BA3a, BA3b), primary motor areas (BA4a [anterior], BA4p [posterior]), premotor area (BA6), and Broca’s area (BA44, BA45). For each region, we extracted average cortical thickness, surface area, volume, and thickness standard deviation, resulting in 36 imaging markers. These markers constituted the structural modality input for the model.In the main analysis, morphometric data from the hemisphere dominant for PD symptoms were used. Statistics of models based on the other hemispheres are presented in the Hemispheric Differences section of the Results.

### 5.4 Data Analysis

All analyses were performed offline using custom-made scripts in MATLAB (MathWorks Inc., Natick, MA, USA).

#### 5.4.1 Neurophysiological Data Preprocessing

Local field potentials within the GPi were recorded in a bipolar configuration using adjacent contacts. All data analyzed were collected during DBS-OFF states where subjects maintained a minimum of at least15 seconds of continuous resting without unnecessary voluntary movements.

Three bipolar ECoG signal pairs were defined:

**Precentral electrodes (Pre-CS):** Contacts immediately anterior to the central sulcus.

**Central electrodes (CS):** Contacts located on either side of the central sulcus.

**Postcentral electrodes (post-CS):** Contacts immediately posterior to the central sulcus (Figure 4B).

Neurophysiological data preprocessing included removal of 60 Hz line noise and discarding segments with electrical or movement artifacts, following previously published procedures [56]. Artifact segments were identified based on criteria such as abnormally high power spectral values, excessive harmonics, and rapid voltage changes. Artifact removal was automated through full-wave rectification, calculation of the first derivative, and application of a five-point median filter. Segments where the first derivative exceeded five standard deviations (SDs) of the subject’s entire recording were considered artifacts and replaced by linear interpolation using data from 2 ms before and after the artifact.

Power spectral density (PSD) was estimated using Thomson’s multitaper method [47] with 1-second consecutive, non-overlapping time windows across a frequency range of 4 to 110 Hz. A frequency bandwidth of ±2 Hz and three tapers were used. To control for inter-subject variability in baseline power, each segmented spectrum was normalized to the total power of the signal for each condition, excluding line noise and its harmonics. Average power within specific frequency bands—alpha (8–11 Hz), low beta (12–20 Hz), high beta (21–35 Hz), and high gamma (65–110 Hz)—was calculated using the multitaper PSDs.

Large matrices containing structural and neurophysiological data were subsequently constructed and prepared for cross-correlation analysis and Sparse Partial Least Squares (SPLS) analysis (Figure 5).Feature Extraction

**Fig. 5:**
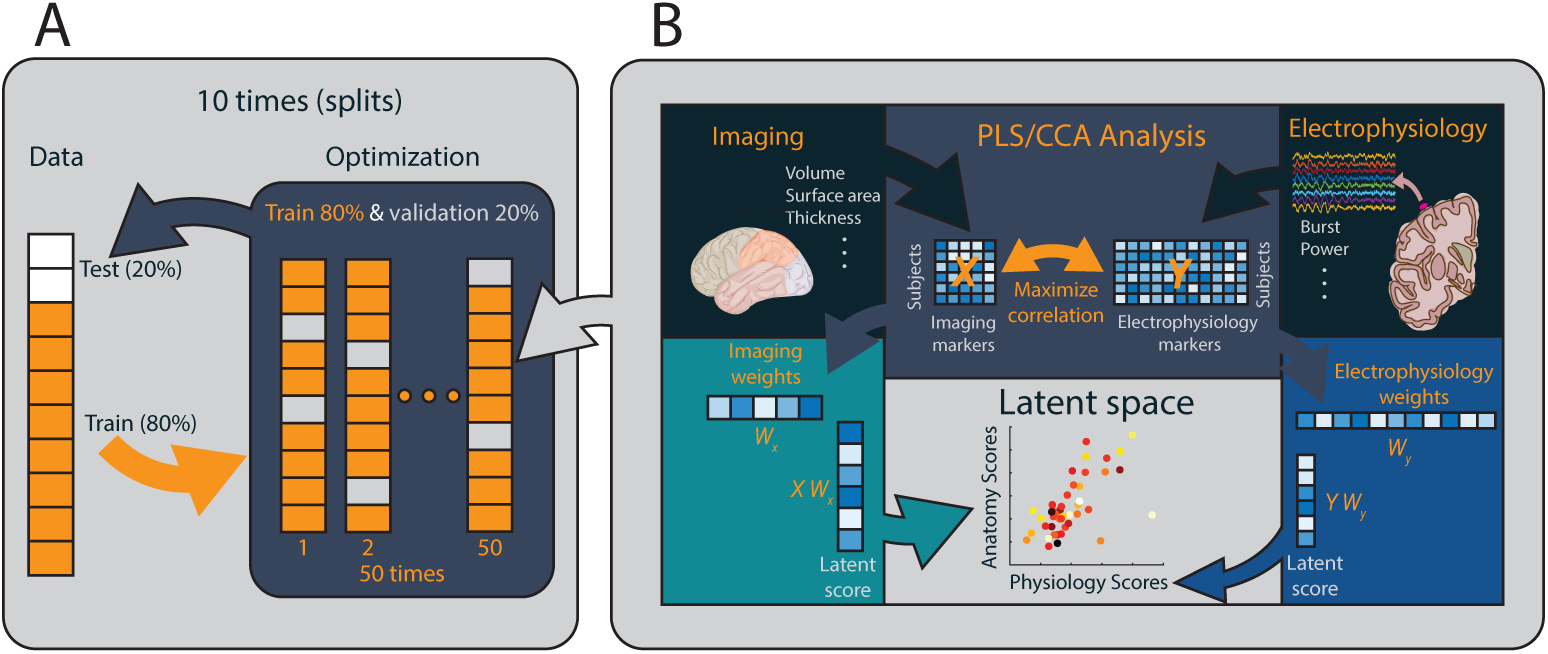
Schematic pipeline of associative correlation analysis. (A) The analysis method illustrated in the figure follows a comprehensive multiple-holdout framework. This approach begins by splitting the original dataset into two parts: an optimization set (comprising 80% of the data) and a holdout set (comprising the remaining 20%) as shown in panel A. The optimization set is divided into a training set (80%) and a validation set (20%). These subsets are used to fit a regularized Partial Least Squares/Canonical Correlation Analysis (PLS/CCA) model and to optimize the regularization parameters through 50 different training and validation iterations. After determining the optimal parameters, the regularized PLS/CCA model is refitted using the entire optimization set and then evaluated on the holdout set with permutation testing to assess its performance. This whole process is repeated 10 times to enhance the robustness and reliability of the model results. (B) The panel B illustrates the PLS/CCA models, which are employed to find weight vectors that maximize the covariance (in the case of PLS) or correlation (in the case of CCA) between linear combinations of brain imaging and behavioral data. The sparse PLS model introduces sparsity constraints, effectively reducing some weights for imaging and electrophysiology variables to zero, thereby selecting the most significant features. The resulting linear combinations or weighted sums of structural and neurophysiological data (derived from matrices *X* and *Y* with corresponding weights Wx and Wy) yield structure and neurophysiology scores (*X* Wx and *Y* Wy) for each participant. These scores are then used to construct an structure-neurophysiology latent space, which represents the relationships between structural and neurophysiological factors across the entire study sample.

Bursting periods were defined as oscillation amplitude periods exceeding the 75th percentile of power for that subject within a specific frequency band and persisting for a minimum of one complete cycle (Figure 1C). We computed the average burst duration, burst amplitude, and burst rate (number of bursts per second) for each subject.To quantify the degree of co-variability between cortico-cortical and cortico-subcortical signal pairs, magnitude-squared coherence was estimated using the multitaper method with the same time window and frequency smoothing parameters as the PSD analysis (±2 Hz frequency bandwidth with three tapers).

For each patient, a total of 92 electrophysiological measures were calculated across five frequency bands and four recording sites (central (CS), precentral (pre-CS), postcentral (Post-CS), and pallidal electrodes). These measures included power, burst duration, burst amplitude, burst rate, cortico-cortical coherence, and corticosubcortical coherence. Cortico-cortical coherences were calculated for three pairs between the central ECoG and the other cortical ECoGs (Pre-CS and Post-CS), and cortico-subcortical coherences were calculated between the pallidal lead with the highest average beta power and the three cortical ECoGs.

#### 5.4.2 Pairwise Cross-Correlation Analysis

We examined the relationships between imaging and neurophysiological measures by calculating Pearson’s correlation coefficients for all possible pairs and organizing the results based on sub-features of each set. Structural measures were divided into two subgroups: Brain regions: Brodmann areas around the general regions where ECoG electrodes were placed, specifically BA1, BA2, BA3a, BA3b, BA4a, BA4p, BA6, BA44, and BA45.

Morphometric parameters: Average surface area, volume, average thickness, Neurophysiological measures were categorized into three subgroups:

Electrode locations: Central, precentral, postcentral, and subcortical sites. Neurophysiological metrics: Power, burst duration, burst amplitude, burst rate, corticocortical coherence (CC), and cortico-subcortical coherence (CS). Frequency bands: Alpha, low beta, high beta, and gamma. We computed the correlation matrix for these subcategories across all 50 subjects using Pearson’s correlation coefficient. All combinations of cross-correlations were visualized using heatmaps, depicting individual correlations (Figure 3) and subgroup averages (Figure 4). To assess the statistical significance of the observed correlations among the 36 variables, we performed permutation tests. The null hypothesis posited that no relationship exists between the variables, and any observed correlation is due to random chance. For the permutation tests, we independently permuted the data for each variable, effectively shuffling the values within each variable to disrupt existing associations while maintaining the original distribution. This process was repeated 1,000 times to generate a distribution of correlation coefficients under the null hypothesis. The p-value for each observed correlation coefficient was calculated as the proportion of permuted correlation coefficients that were as extreme or more extreme than the observed value:

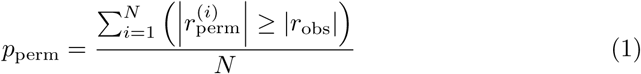

where *p*_perm_ represents the p-value obtained from the permutation tests, *i* denotes the permutation iteration, 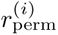 is the correlation coefficient from the permuted data, *r*_obs_ is the observed correlation coefficient from the original dataset.

To adjust for multiple comparisons, we employed the False Discovery Rate (FDR) control using the Benjamini-Hochberg procedure. The permutation p-values were compared against thresholds determined by:

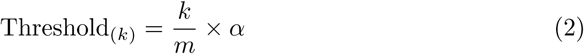

Where *k* is the rank of the p-value after sorting in ascending order, *m* is the total number of tests performed in the category, and *α* is the desired FDR, set to 0.05 in this study. We rejected all null hypotheses for which the p-values were less than or equal to their corresponding threshold.

#### 5.4.3 Sparse Partial Least Squares Analysis

Recognizing that pairwise analysis does not account for associative effects among variables, we employed Sparse Partial Least Squares (SPLS) regression to explore multivariate associations between structural and electrophysiological measures [48, 49]. SPLS is well-suited for capturing complex relationships between two data modalities, especially in high-dimensional datasets. We utilized the SPLS implementation developed by 50 to train, regularize, and evaluate the model. SPLS optimizes the covariance between linear combinations of imaging and neurophysiological variables, producing weight vectors that indicate the contribution of each variable to the identified associations. The method includes sparsity constraints, enhancing model interpretability by selecting a subset of important features and focusing on the most relevant variables.

#### 5.4.4 Model Training, Optimization, and Validation

To ensure generalizability and avoid overfitting, we employed a multiple holdout framework for optimizing regularization parameters and evaluating the SPLS models (Figure 2). The SPLS model was trained on an optimization set comprising 80% of the data, and the identified multivariate associations were assessed on a separate holdout set comprising the remaining 20%. The optimization set was further divided into training and validation subsets to fine-tune regularization parameters. Optimal parameters were selected based on the ability to enhance generalization performance, determined by the out-of-sample correlation on the validation set, using a combined criterion of stability and generalizability. Generalizability was quantified as the average out-of-sample correlation on the validation and holdout sets. Stability was assessed by calculating the average similarity of the weight vectors (corrected overlap for SPLS) across different data splits. To validate the robustness of the SPLS model, the entire process was repeated 10 times, following methodologies from previous studies [50–53]. This iterative approach allowed for the identification of structure-neurophysiology associations that are both stable and generalizable to new data. To identify symptom-related factors using UPDRS scores, we performed stepwise multiple regression analyses using anatomical and physiological biomarkers weighted by SPLS-derived split weights. The stepwise approach was optimized using the Akaike Information Criterion (AIC) to select the best subset of predictors. Separate models were created for anatomical and physiological variables, and the analysis was conducted to determine which biomarkers significantly contributed to explaining UPDRS scores.

#### 5.4.5 Essential Tremor (ET) Cohort Analysis

To evaluate whether the SPLS/CCA model and latent space were specific to PD pathophysiology, we tested the model on a cohort of 24 essential tremor (ET) patients. Data collection and the calculation of morphometric and neurophysiological variables were conducted in the same manner as for the main PD cohort. The SPLS weights from the best-performing split identified in the PD analysis were then applied to calculate the latent space for the ET patients. The correlation between anatomical and physiological latent scores was computed to determine whether the model captured meaningful associations in the ET cohort.

## 5.5 Data availability

The datasets analyzed during the current study are available in the Brain Integrated Resource for Human Anatomy and Intracranial Neurophysiology(B(RAIN2)) repository, https://doi.org/10.18120/zmah-2816.

## 5.6 Code availability

The underlying code for this study is not publicly available but may be made available to qualified researchers on reasonable request from the corresponding author.

## 5.7 Acknowledgment

We gratefully thank Sahil Chilukur for his dedicated support in patient recruitment and data collection. This work has been supported by the National Institute of Neurological Disease and Stroke of the National Institutes of Health (https://www.ninds.nih.gov/) under award number R01NS097782 and RF1MH121373. The funder played no role in study design, data collection, analysis and interpretation of data, or the writing of this manuscript.

## 5.8 Author contributions

KM, AA, and NP: experimental design and setup, data acquisition, analysis design, data interpretation, and manuscript writing. KM: implementing the data analysis code and algorithms. AA: Imaging data analysis and electrode localization. NP: Surgical procedures. All authors read and approved the final manuscript.

## 5.10 Declaration on the Use of Artificial Intelligence

During the preparation of this work, the authors used AI-powered language tools for assistance with spelling and grammar. All AI-generated suggestions were manually reviewed and edited by the authors to ensure accuracy and clarity. The authors take full responsibility for the content of this publication.

## References

[1] Brown, P. et al. Dopamine dependency of oscillations between subthalamic nucleus and pallidum in Parkinson’s disease. J. Neurosci. 21, 1033–1038 (2001).

[2] Malekmohammadi, M., AuYong, N., Ricks-Oddie, J., Bordelon, Y., Pouratian, N. Pallidal deep brain stimulation modulates excessive cortical high beta phase amplitude coupling in Parkinson disease. Brain Stimulat. 11, 607–617 (2018).

[3] Neumann, W.-J. et al. Subthalamic synchronized oscillatory activity correlates with motor impairment in patients with Parkinson’s disease. Mov. Disord. Off. J. Mov. Disord. Soc. 31, 1748–1751 (2016).

[4] Oswal, A. et al. Deep brain stimulation modulates synchrony within spatially and spectrally distinct resting state networks in Parkinson’s disease. Brain 139, 1482–1496 (2016).

[5] Hirschmann, J. et al. Longitudinal Recordings Reveal Transient Increase of Alpha/Low-Beta Power in the Subthalamic Nucleus Associated With the Onset of Parkinsonian Rest Tremor. Front. Neurol. 10, 145 (2019).

[6] Little, S., Pogosyan, A., Kuhn, A. A., Brown, P. Beta band stability over time correlates with Parkinsonian rigidity and bradykinesia. Exp. Neurol. 236, 383–388 (2012).

[7] Malekmohammadi, M. et al. Pallidal stimulation in Parkinson disease differentially modulates local and network beta activity. J. Neural Eng. 15, (2018).

[8] Van Wijk, B. C. M. et al. Low-beta cortico-pallidal coherence decreases during movement and correlates with overall reaction time. NeuroImage 159, 1–8 (2017).

[9] Lofredi, R. et al. Beta bursts during continuous movements accompany the velocity decrement in Parkinson’s disease patients. Neurobiol. Dis. 127, 462–471 (2019).

[10] O’Keeffe, A. B., Malekmohammadi, M., Sparks, H., Pouratian, N. Synchrony Drives Motor Cortex Beta Bursting, Waveform Dynamics, and Phase-Amplitude Coupling in Parkinson’s Disease. J. Neurosci. 40, 5833–5846 (2020).

[11] Tinkhauser, G. et al. Beta burst dynamics in Parkinson’s disease off and on dopaminergic medication. Brain 140, 2968–2981 (2017).

[12] Pereira, J. B. et al. Initial cognitive decline is associated with cortical thinning in early Parkinson disease. Neurology 82, 2017–2025 (2014).

[13] Zhang, L. et al. Cortical Thinning and Cognitive Impairment in Parkinson’s Disease without Dementia. IEEE/ACM Trans. Comput. Biol. Bioinform. 15, 570–580 (2018).

[14] Gao, Y. et al. Changes in Cortical Thickness in Patients With Early Parkinson’s Disease at Different Hoehn and Yahr Stages. Front. Hum. Neurosci. 12, 469–469 (2018).

[15] Segura, B. et al. Cortical thinning associated with mild cognitive impairment in Parkinson’s disease. Mov. Disord. 29, 1495–1503 (2014).

[16] Jia, X. et al. Longitudinal Study of Gray Matter Changes in Parkinson Disease. *Am*. J. Neuroradiol. 36, 2219–2226 (2015).

[17] Nürnberger, L., et al. Longitudinal changes of cortical microstructure in Parkinson’s disease assessed with T1 relaxometry. NeuroImage Clin. 13, 405–414 (2017).

[18] Tessa, C. et al. Progression of brain atrophy in the early stages of Parkinson’s disease: A longitudinal tensor-based morphometry study in de novo patients without cognitive impairment. Hum. Brain Mapp. 35, 3932–3944 (2014).

[19] Brenneis, C. et al. Voxel-based morphometry detects cortical atrophy in the Parkinson variant of multiple system atrophy. Mov. Disord. 18, 1132–1138 (2003).

[20] Weintraub, D. Neurodegeneration Across Stages of Cognitive Decline in Parkinson Disease. Arch. Neurol. 68, 1562 (2011).

[21] Burton, E. J. Cerebral atrophy in Parkinson’s disease with and without dementia: a comparison with Alzheimer’s disease, dementia with Lewy bodies and controls. Brain 127, 791–800 (2004).

[22] Morgen, K. et al. Structural Brain Abnormalities in Patients with Parkinson Disease: A Comparative Voxel-Based Analysis Using T1-Weighted MR Imaging and Magnetization Transfer Imaging. *Am*. J. Neuroradiol. 32, 2080–2086 (2011).

[23] Nagano-Saito, A. et al. Cerebral atrophy and its relation to cognitive impairment in Parkinson disease. Neurology 64, 224–229 (2005).

[24] Summerfield, C. et al. Structural Brain Changes in Parkinson Disease With Dementia: A Voxel-Based Morphometry Study. Arch. Neurol. 62, 281 (2005).

[25] Cordato, N. J., Duggins, A. J., Halliday, G. M., Morris, J. G. L., Pantelis, C. Clinical deficits correlate with regional cerebral atrophy in progressive supranuclear palsy. Brain 128, 1259–1266 (2005).

[26] Sterling, N. W. et al. Structural Imaging and Parkinson’s Disease: Moving Toward Quantitative Markers of Disease Progression. J. Park. Dis. 6, 557–567 (2016).

[27] Sanmartino, F. et al. Subthalamic Beta Activity in Parkinson’s Disease May Be Linked to Dorsal Striatum Gray Matter Volume and Prefrontal Cortical Thickness: A Pilot Study. Front. Neurol. 13, 799696 (2022).

[28] Cohen, S. L., Woo Choi, J., Toga, A. W., Pouratian, N., Duncan, D. Exaggerated High-Beta Oscillations are Associated with Cortical Thinning at the Motor Cortex in Parkinson’s Disease. In: 2023 45th Annual International Conference of the IEEE Engineering in Medicine & Biology Society (EMBC), 1–4. IEEE, Sydney, Australia, 2023. doi:10.1109/EMBC40787.2023.10341040.

[29] Ibarretxe-Bilbao, N. et al. Progression of cortical thinning in early Parkinson’s disease. Mov. Disord. 27, 1746–1753 (2012).

[30] Sarasso, E., Agosta, F., Piramide, N., Filippi, M. Progression of grey and white matter brain damage in Parkinson’s disease: a critical review of structural MRI literature. J. Neurol. 268, 3144–3179 (2021).

[31] Park, C. et al. Simulating the progression of brain structural alterations in Parkinson’s disease. npj Park. Dis. 8, 86 (2022).

[32] Wilson, H., Niccolini, F., Pellicano, C., Politis, M. Cortical thinning across Parkinson’s disease stages and clinical correlates. J. Neurol. Sci. 398, 31–38 (2019).

[33] Pereira, J. B. et al. Assessment of cortical degeneration in patients with Parkinson’s disease by voxel-based morphometry, cortical folding, and cortical thickness. Hum. Brain Mapp. 33, 2521–2534 (2012).

[34] Chen, Z., Li, G., Zhou, L., Zhang, L., Liu, J. Altered structural-functional coupling in Parkinson’s disease. Preprint at 10.1101/2023.01.18.23284750 (2023).

[35] Zarkali, A. et al. Organisational and neuromodulatory underpinnings of structural-functional connectivity decoupling in patients with Parkinson’s disease. *Commun*. Biol. 4, 86 (2021).

[36] Pollok, B. et al. Increased SMA–M1 coherence in Parkinson’s disease — Pathophysiology or compensation? Exp. Neurol. 247, 178–181 (2013).

[37] Droby, A. et al. The interplay between structural and functional connectivity in early stage Parkinson’s disease patients. J. Neurol. Sci. 442, 120452 (2022).

[38] Wilson, H. et al. Cortical thinning across Parkinson’s disease stages and clinical correlates. J. Neurol. Sci. 398, 31–38 (2019).

[39] Pfurtscheller, G., Stancák, A., Neuper, Ch. Event-related synchronization (ERS) in the alpha band — an electrophysiological correlate of cortical idling: A review. Int. J. Psychophysiol. 24, 39–46 (1996).

[40] Brown, P. Abnormal oscillatory synchronisation in the motor system leads to impaired movement. Curr. Opin. Neurobiol. 17, 656–664 (2007).

[41] Conte, A., Khan, N., Defazio, G., Rothwell, J. C., Berardelli, A. Pathophysiology of somatosensory abnormalities in Parkinson disease. Nat. Rev. Neurol. 9, 687–697 (2013).

[42] Wang, Y. et al. Hypo-connectivity of the primary somatosensory cortex in Parkinson’s disease: a resting-state functional MRI study. Front. Neurol. 15, 1361063 (2024).

[43] Perinelli, A., Castelluzzo, M., Tabarelli, D., Mazza, V., Ricci, L. Relationship between mutual information and cross-correlation time scale of observability as measures of connectivity strength. Chaos Interdiscip. J. Nonlinear Sci. 31, 073106 (2021).

[44] Bergamino, M. et al. Structural connectivity and brain network analyses in Parkinson’s disease: A cross-sectional and longitudinal study. Front. Neurol. 14, 1137780 (2023).

[45] Wiesman, A. I. et al. Alterations of Cortical Structure and Neurophysiology in Parkinson’s Disease Are Aligned with Neurochemical Systems. Ann. Neurol. 95, 802–816 (2024).

[46] Randazzo, M. J. et al. Three-dimensional localization of cortical electrodes in deep brain stimulation surgery from intraoperative fluoroscopy. NeuroImage 125, 515–521 (2016).

[47] Thomson, D. J. Spectrum estimation and harmonic analysis. Proc. IEEE 70, 1055–1096 (1982).

[48] Abdi, H. Partial least squares regression and projection on latent structure regression (PLS Regression). WIREs Comput. Stat. 2, 97–106 (2010).

[49] Wold, H. Partial Least Squares. In: Encyclopedia of Statistical Sciences (eds. Kotz, S., Read, C. B., Balakrishnan, N., Vidakovic, B.) (Wiley, 2004). doi:10.1002/0471667196.ess1914.

[50] Mihalik, A. et al. Canonical Correlation Analysis and Partial Least Squares for Identifying Brain–Behavior Associations: A Tutorial and a Comparative Study. Biol. Psychiatry Cogn. Neurosci. Neuroimaging 7, 1055–1067 (2022).

[51] Baldassarre, L., Pontil, M., Mouräo-Miranda, J. Sparsity Is Better with Stability: Combining Accuracy and Stability for Model Selection in Brain Decoding. Front. Neurosci. 11, (2017).

[52] Mihalik, A. et al. Brain-behaviour modes of covariation in healthy and clinically depressed young people. Sci. Rep. 9, 11536 (2019).

[53] Monteiro, J. M., Rao, A., Shawe-Taylor, J., Mouräo-Miranda, J. A multiple holdout framework for Sparse Partial Least Squares. J. Neurosci. Methods 271, 182–194 (2016).

[54] Kühn, A. A., et al. Event-related beta desynchronization in human subthalamic nucleus correlates with motor performance. Brain 127, 735–746 (2004).

[55] Litvak, V. et al. Movement-related changes in local and long-range synchronization in parkinson’s disease revealed by simultaneous magnetoencephalography and intracranial recordings. J. Neurosci. 32, 10541–10553 (2012).

[56] AuYong, N., Malekmohammadi, M., Ricks-Oddie, J. Pouratian, N. Movement-modulation of local power and phase amplitude coupling in bilateral globus pallidus interna in Parkinson disease. Front. Hum. Neurosci. 12, 1–13 (2018).

